# An evolutionarily conserved microRNA, miR-185-5p, regulates key pathways that may contribute to implantation failure

**DOI:** 10.1101/2025.01.15.633123

**Authors:** William Smith, Jessica C. Edge, Haidee Tinning, Zenab Butt, Fani Deligianni, Cintli Morales, Joanne Muter, Jan Brosens, Mariano Mascarenhas, Harish Bhandari, Mary J. O’Connell, Emma Lucas, Nigel Simpson, Niamh Forde

## Abstract

Recurrent implantation failure (RIF) is defined after three or more good quality embryo transfers following *in vitro* fertilisation without a successful pregnancy outcome. Many factors contribute to RIF however, the endometrial contribution remains unclear. Previous work by our group has identified a micro-RNA (miRNA) miR-185-5p as conserved across placental mammal irrespective of implantation strategies. We tested the hypothesis that miR-185-5p and the pathways it regulates, may be disrupted in the endometria of women with RIF. A human endometrial epithelial cells line (Ishikawa cells) was transfected with mimics or inhibitors for miR-185-5p for 24 (for implantation assay) or 48 hr (for proteomic analysis) along with non-targeting controls. There was a significant different in percentage attachment of BeWoW spheroids to cells transfected with miR-185-5p mimic compared to inhibitor (P<0.05). Transfection of epithelial cells with miR-185-5p altered expression of 1450 (mimic alone) and 509 (inhibitor alone) proteins respective of which, 146 were modified by both. Comparison of predicted targets of miR-185-5p, proteins modified by this study, and key endometrial genes from the literature determined genes and proteins associated with CNP family were further investigated in biopsies from individuals with (n=10) and without RIF (n=9) with *CSNK1D* expression significantly lower (p<0.05) in individuals with RIF. Collectively these data demonstrate that miR-185-5p modifies pathways that are important for successful implantation in humans.

## INTRODUCTION

Recurrent implantation failure (RIF) is diagnosed typically after three or more good quality embryo transfers following *in vitro* fertilisation (Polanski et al., 2014; Somigliana et al., 2018; Mascarenhas et al., 2022; Cimadomo et al., 2023; Pirtea et al., 2023). Although there are global variations in the definition of RIF, it is believed to affect < 5% of couples with infertility (Pirtea et al., 2021; Pirtea et al., 2023). Although a range of overlapping routine investigations are typically considered in patients with RIF and early pregnancy loss, no specific and reliable markers of endometrial dysfunction have been accepted for routine clinical application RIF patients (Aplin and Stevens, 2022; HFEA, 2023; Cimadomo et al., 2023). More recently, ‘omics technologies have begun to identify themes among key molecules, pathways, and genes that either contribute to, or are highly associated with, changes in the molecular phenotypes in the endometrium (Lessey and Arnold, 1998; Dahmoun et al., 1999; Giudice, 2004; Talbi et al., 2006; Achache and Revel, 2006; Gellersen and Brosens, 2014; Koot et al., 2016; Aplin and Ruane, 2017). These include key transition points during the menstrual cycle, reflecting the synchronous response to ovarian sex-steroid hormones (Emera et al., 2012; Alvergne and Högqvist Tabor, 2018) and have been explored in endometrial biopsies from women with normal menstrual cycles (Kao et al., 2002; Talbi et al., 2006). There is also a surge of interest in the molecular phenotypes within the endometrium, relating to changes in local inflammatory response, including alterations in the behaviour of specific leukocyte subpopulations - the uterine natural killer cells (uNKs) (Kuroda et al., 2013; Chong and Quenby, 2016). These, along with populations of local and circulating progenitor cells, are key to regenerating the endometrium on a cyclical basis, including differentiation of decidualised stromal cells to facilitate embryo recognition and pregnancy tolerance (Ashkar et al., 2003; van den Heuvel et al., 2005; Xiong et al., 2013; Gellersen and Brosens, 2014; Brighton et al., 2017; Lucas et al., 2020).

The role of endometrial luminal epithelial cells during early embryo apposition was highlighted almost thirty years ago by Glasser and Mulholland (Glasser and Mulholland, 1993). Whilst appropriately nourished embryos have been shown to attach to any surface *in vitro* (Enders et al., 1981; Sengupta et al., 1986), the discriminatory events that occur *in vivo* are yet to be elucidated. Early embryo-endometrial interactions have been evaluated more recently in mice (Salker et al., 2011) and ruminants (Spencer et al., 2004), however studies in humans remain limited from both a medical ethics perspective and because of a lower embryo yield (Singh et al., 2010). Nevertheless, there are similarities across species in some of the signalling mechanisms around receptivity to implantation (Psychoyos, 1986; Enders, 2000; Singh et al., 2010). Expression of specific cell surface molecules and ligands has previously been shown to be hormonally regulated during the window of implantation (Apparao et al., 2001; Gellersen and Brosens, 2014) and defective expression of these molecules may contribute to reproductive failure phenotypes. Decidualisation (a process that is not observed in all mammalian species) is associated with a transcriptomic shift, contributing to the regulation of transcription factors, and secretion of growth factors and cytokines, with widespread endocrine and paracrine signalling, changes in adhesion molecules and extracellular matrix (ECM) ligands, and the production of other anti-inflammatory and pro-inflammatory molecules (Giudice, 2004; Garrido-Gomez et al., 2011; Gellersen and Brosens, 2014; Altmäe et al., 2017). These require a tightly regulated dialogue between the endometrial leukocyte subpopulations (including UNKs, regulatory T-cells, macrophages and uterine dendritic cells, or uDCs) and transformed decidualising endometrial stromal cells (dESCs) (van Mourik et al., 2009; Gargett et al., 2009; Kuroda et al., 2013; Robertson et al., 2018; Liu et al., 2020), contributing to endometrial plasticity and the potential to adapt to embryonic (Capalbo et al., 2016; Gross et al., 2017; Kong et al., 2021) and paternal signals (Moldenhauer et al., 2009; Robertson et al., 2009; Robertson and Sharkey, 2016) with co-regulation of cell-fate decisions appearing to be critical during cyclical remodelling of the endometrium (Giudice, 2004; Gellersen and Brosens, 2014; Brighton et al., 2017; Kong et al., 2021). Clearly there are many factors that regulate the transcriptional and translational landscape but the upstream control of these is largely unknown.

One set of molecules that may regulate endometrial function/ dysfunction are microRNAs. The association between pathophysiology and dysregulated miRNAs has been widely published, with miRNAs being employed as biomarkers of disease, and demonstrating both oncogenic and tumour-suppressive potential (Hammond, 2007; Cortez et al., 2011; Hayes et al., 2014; Peng and Croce, 2016; Rupaimoole and Slack, 2017). In contrast, a significant number of miRNAs have also been associated with regulation of normal physiological processes, including innate and adaptive immune responses through regulation of haematopoietic stem cells (O’Connell et al., 2010). Previous work from our group has identified several conceptus derived proteins that are highly conserved amongst placental mammals and have been shown to regulate a core set of placental mammal-conserved miRNAs in species with different implantation strategies (Forde et al., 2015; Taylor et al., 2023), one of which is miR-185-5p. Given the conserved nature of this miRNA across mammal species with different implantation strategies, as well as its function in other epithelial cell systems, we tested the hypothesis that miR-185-5p and the pathways it regulates, may be disrupted in the endometria of women with RIF.

## MATERIALS AND METHODS

Unless otherwise stated all consumables were sourced from Sigma Aldrich (UK).

### *In vitro* implantation assay

#### Cell transfections

Human endometrial epithelial cells (*Ishikawa* cells: ECACC - #99040201) cultured at 37°C and 5% CO2 (37°C/5% CO2) were maintained in 50:50 Gibco® Dulbecco’s Modified Eagle Medium: Ham’s Nutrient Mixture F-12 (DMEM/F-12) (ThermoFisher Scientific, USA) with 10% FBS Gold (ThermoFisher Scientific, USA) and 1% Gibco® gentamicin-streptomicin-penicillin (GSP). Prior to transfection, passage 37 cells were changed to antibiotic-free media with serum. Cells were seeded at a density of 130,000 cells/mL in 24-well plates (n=3 biological replicates, n=2 technical replicates), media aspirated, and cells washed twice with PBS. The following media and treatments were then added (n=6 wells per treatment): **1) Control**: 500μL media (antibiotic-free media DMEM/F-12 + 10% FBS Gold), **2) Vehicle Control**: 400μL media + 100μL DharmaFECT 2 solution (50μL OptiMEM without the addition of mimic or inhibitor combined with 1μL DharmaFECT 2 in 50μL OptiMEM), **3) Non-Targeting Mimic**: 400μL media + DharmaFECT 2 solution (with 0.2μL [100µM] miRIDIAN non-targeting mimic; NTM), **4) Non-Targeting Inhibitor**: 400μL media + DharmaFECT 2 solution (with 0.2μL [100µM] miRIDIAN non-targeting inhibitor; NTI), **5) miR-185-5p Mimic**: 400μL media + DharmaFECT 2 solution (with 0.2μL [100µM] miRIDIAN miR-185-5p mimic; 185-M), **6) miR-185-5p Inhibitor**: 400μL media + DharmaFECT 2 solution (with 0.2μL [100µM] miRIDIAN miR-185-5p inhibitor; 185-I). All 24-well plates were incubated for 48 hr. To confirm miR-185-5p expression, all wells in one column per 24-well plate were treated with 0.025% trypsin (incubated for 3 mins followed by 5mL complete media to inactivate trypsin), transferred to separate sterile 1.5mL tubes, centrifuged at 500 × g for 5 mins, media aspirated and 700μL QIAzol lysis reagent (Qiagen®) added to each tube and samples snap frozen in liquid nitrogen and stored at −80°C. The remaining wells were used to evaluate spheroid-epithelial attachment (see below).

#### BeWo spheroid formation and implantation assay

BeWo Cells (ECACC:#86082803) were cultured in antibiotic-free and serum-free media (DMEM-F12). Once >50% confluent, the T75 flasks of BeWo cells were then lifted using 0.025% trypsin for 3 min at 37°C (followed by inactivation of trypsin with complete media). The wells of a 6-well plate were coated with anti-adhesive polymer, polyvinylpyrrolidone (PVP), for 30 mins, followed by PBS to avoid drying-out. The lifted cells were transferred to a 15mL Falcon tube and centrifuged at 500 × g for 5 mins, the supernatant aspirated and then washed in 5mL PBS. Tubes were then centrifuged at 500 × g for 5 mins, the supernatant aspirated and pellet resuspended again in antibiotic-free *and* serum-free media (DMEM/F-12). At least 500,000 cells/well were transferred to each well of the 6-well plate (2 × T75 flask producing enough BeWo cells for 3 wells). The BeWo cells were then incubated (37°C/5% CO_2_) on an orbital shaker for 4 hr (60 rpm), followed by gently pipetting up and down of the contents of each well to disrupt any large BeWo clumps. The 6-well plate of BeWo cells were then incubated overnight, although subsequently not on the orbital shaker (37°C/5% CO_2_) to achieve spheroid formation. Spheroids equivalent in size to an implanting blastocyst (approximately 50-100μm in diameter), were isolated using appropriate cell strainers. Briefly, the BeWo cells were rinsed with antibiotic-free and serum-free media and passed through a 100 μm cell strainer positioned on top of a 40 μm cell strainer. The 40μm strainer was then inverted over a new 50mL Falcon tube and rinsed again to elute the spheroids. The eluted spheroids and treated hEECs were transferred immediately (separately) to the EVOS cell imaging system (ThermoFisher Scientific). Next, 100μL of eluted spheroids with a minimum of 50 spheroids present were added to each of the treatments outlines above. Distribution of spheroids across each well was achieved by gently rocking the plates with a cross-pattern movement. Each plate was then incubated for 1hr (37°C/5% CO2). Spheroid-epithelial attachment was then evaluated as described below.

To confirm appropriate transfection and changes to miR-185-5p expression, total RNA was extracted from hEECs following the 48hr incubation step, and prior to Be-Wo co-culture (one column from each 24-well plate). RNA exaction was performed using the Qiagen® miRNeasy® Mini Kit with on-column DNase digestion according to manufacturer’s instructions. Following RNA extraction all samples were prepared for reverse transcription using the miRCURY LNA RT Kit (Qiagen®, UK) and reverse transcribed to cDNA using the corresponding manufacturer’s instructions as previously described (Hume et al., 2023). Confirmation of miR-185-5p expression was evaluated using the Qiagen® miRCURY LNA PCR Assay for miR-185-5p as per manufacturer’s instructions. Analysis of miR-185-5p expression was completed using the Roche LightCycler®96 according to manufacturer’s instructions.

#### EVOS Cell Imaging and implantation analysis

Following initial co-incubation (37°C/5% CO2) for 30 min each 24-well plate was photographed at 4× magnification using the EVOS Cell Imaging System (from ThermoFisher Scientific), with adjusted phase contrast (with manually adjusted focus), imaging the entirety of each well (scan area set to 100%), with sequential images of each well ‘stitched-together’. Once all images were obtained, media from each well was carefully aspirated (avoiding contact with the centre of the well) and 500μL 10% formalin added to each well and the plate incubated at RT for 20 mins. The formalin was then aspirated, each well washed with 500μL PBS and all media aspirated. All plates were then re-photographed using the same method as before. Once all images were obtained (pre- and post-application of formalin), ImageJ (https://imagej.nih.gov/ij/download.html) was used to count spheroids in each well to determine spheroid-epithelial attachment percentage. To ensure effective and systematic counting of spheroids, before and after images obtained from EVOS for each well were viewed (and BeWo spheroids counted) simultaneously, therefore helping to avoid potentially miscounting or over-representing artefacts in the wells. Mean spheroid-epithelial attachment percentage was then compared between treatment groups of interest and any significant differences confirmed with GraphPad Prism (Version 9.5.1) using an unpaired, two-tailed *t*-test (α = 0.05). A one-way analysis of variance (ANOVA) with Dunnett’s multiple comparisons (with multiplicity adjusted p values (Staffa and Zurakowski, 2020) was also used to assess differences across all treatment groups (relative to vehicle control). Individual spheroid-epithelial attachment percentage was demonstrated using modified scatter plots, with mean spheroid epithelial attachment and 95% CI plotted. The 95% CIs were calculated from the SEM and adjusted for n-1 degrees of freedom given the small number of samples (Campbell and Swinscow, 2002).

### Proteomic analysis of miR-185-5p targets in endometrial epithelial cells

#### Cell culture and proteomics

Ishikawa cells were grown as described above and seeded in at a density of 200,000 cells/mL in 2mL wells. Cells were treated according to groups 1-5, and 7 as above for 48-hr, pellets of cells snap frozen prior to protein extraction. To investigate altered protein abundance in response to miR-185-5p, samples were processed for tandem mass tag (TMT) using 11-Plex reagents at the Bristol Core Proteomics Facility, University of Bristol as previously described (De Bem et al., 2021; Hume et al., 2023). Briefly, 50μg of protein was digested with 1.25 μg trypsin overnight and labelled with 11-Plex reagents in line with manufacturer’s protocol (ThermoFisher Scientific). The labelled samples were then pooled and 100 μg of the pooled sample was desalted using a SepPak cartridge (Waters Corporation) as per manufacturer’s protocol. The eluate was evaporated to dryness and resuspended in 20mM ammonium hydroxide (pH 10) and underwent high pH reverse-phase chromatography using the Ultimate 3000 liquid chromatography system (Thermo Scientific). Using an XBridge BEH C18 column (Waters Corporation, UK), peptides could be shown to elute at different gradients of 20mM ammonium hydroxide in acetonitrile (pH 10), ranging from 0 to 95% in 60 mins. The fractions identified were evaporated to dryness and then resuspended in 1% formic acid prior to nanoscale liquid chromatography mass spectrometry (nano-LC-MS) using Orbitrap Fusion Lumos mass spectrometer (Thermo Scientific). Peptides were ionized by nano-electrospray ionisation (2.0kV) using a stainless-steel emitter with a 30μm diameter and capillary temperature of 300°C. All spectra were acquired using Xcalibur 3.0 software (Thermo Scientific). Raw data from nano-LC-MS / TMT-spectrometry were processed and quantified using Proteome Discoverer software version 2.1 (Thermo Scientific). Master protein selection was enhanced by in-house script prioritising UniProt annotations in protein candidates with equal identification or quantification metrics. The mass spectrometry data were searched against the human UniProt database retrieved 14-10-2021 and updated with additional annotations on 15-11-2021.

#### Bioinformatics analysis of proteomics data and downstream analysis

Protein abundances were Log_2_ transformed to approximate Normality of data distribution. Statistically significant differences in protein abundance between samples was determined using the paired *t*-test and false discovery rate (FDR) corrected using the Benjamin-Hochberg method. Principal component analyses (PCA) was performed using FactoMineR package and plotted using the ggplot2 package using statistical analysis tool R. PCA were produced to measure variation between samples, with a potential range of factors contributing to variation during sample analysis. Volcano plots for comparisons were created using -log10 p-value of each protein against the log2 fold change. Proteins where p < 0.05 were highlighted in yellow, and proteins where p < 0.05 and Log_2_ fold change (Log2FC) > 1, or where p < 0.05 and Log2FC < −1 were highlighted in red.

Data were filtered and quality controlled, producing a data set containing a list of 9734 proteins or protein fragments (including duplicates for some proteins). This complete data set from subsequently evaluated by systematically comparing each treatment group (control, lipofectamine [vehicle control], non-targeting mimic, non-targeting inhibitor, miR-185-5p mimic, miR-185-5p inhibitor) against other treatment groups (e.g., control vs lipofectamine, miR-185-5p mimic vs NTM etc.) selecting out only statistically significant differences in protein abundance. Each comparison created a list of significantly abundant proteins (*t*-test with a *p* value < 0.05). While the use of false discovery rate (FDR) is a more stringent test of protein abundance, it was deemed too restrictive and potentially would have excluded proteins of interest. Once a list of proteins was generated for each treatment group, the lists were cleared of duplicates, and numbers of proteins within each list were then compared against each other using Venn diagram software Venny 2.1 (https://bioinfogp.cnb.csic.es/tools/venny/), identifying common and unique proteins (Schematic of data analysis in Figure 1).

**Figure 1.**
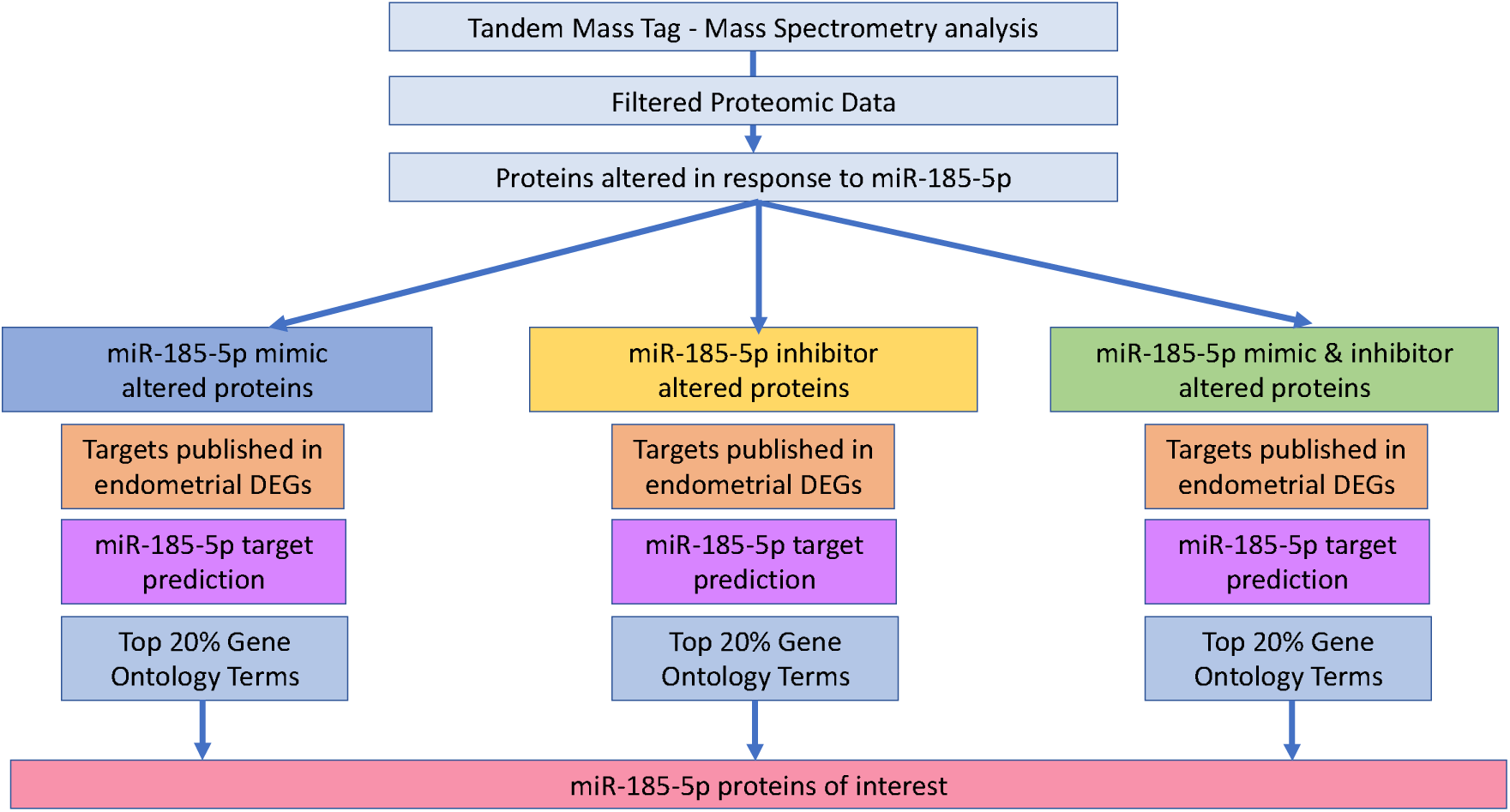
Flow chart illustrating the systematic filtering of proteomic analysis data and identification of miR-185-5p proteins of interest. Tandem-mass-tag (TMT) spectrometry data from Bristol Core Proteomics Facility (BCPF) was filtered to identified statistically significant differences in protein abundance in each treatment group (C – control, L – lipofectamine or vehicle control, NTM – non-targeting mimic, NTI – non-targeting inhibitor, 185-M - miR-185-5p mimic, and 185-I – miR-185-5p inhibitor). Significant differences in protein abundance were compared between treatment groups, identifying a list of proteins specific to miR-185-5p mimic or inhibitor (with a small number of proteins found in both). Each of these protein lists were investigated to determine their functional enrichment in pathways of interest using online functional enrichment toolkits WebGestalt and g:Profiler, revealing functionally enriched gene ontology (GO) terms related to these proteins. In addition to this, targets identified from the BCPF dataset were compared against online miRNA target prediction (miRDB, miRTarBase and TargetScan) and published differentially expressed endometrial genes. Targets found to be significantly represented in each group were selected out for further analysis in RIF clinical samples.

### Comparison of miR-185-5p *in vitro* produced targets to publicly available data sets

#### Comparison to computationally predicted targets

We performed a cross-comparison of the targets predicted computationally for miR-185-5p from miRDB (http://mirdb.org/), miRTarBase (https://mirtarbase.cuhk.edu.cn/) and TargetScan (https://www.targetscan.org/vert_80/) with targets identified *in vitro*.

#### Published differentially expressed endometrial genes (2002-2019)

Web-based searches using PubMed (https://pubmed.ncbi.nlm.nih.gov/) identified thirty-five studies which investigated endometrial gene expression with respect to functional (and dysfunctional) changes across different mammalian species (Table 1). Human gene orthologs were identified from non-human protein accession numbers (and other identifiers such as corresponding NCBI reference sequence) using UniProt (https://www.uniprot.org/), National Institute of Health’s National Centre for Biotechnology Information (https://www.ncbi.nlm.nih.gov/gene/), HUGO Gene Nomenclature (https://www.genenames.org/), Online Mendelian Inheritance in Man (OMIM; https://omim.org/) and Ensembl (https://www.ensembl.org/index.html). This list of published differentially expressed genes (DEGs) was used to draw out proteins of interest from the filtered proteomic data. Additional cross-comparisons could also be made against predicted and functionally-enriched targets (see below).

**Table 1.**
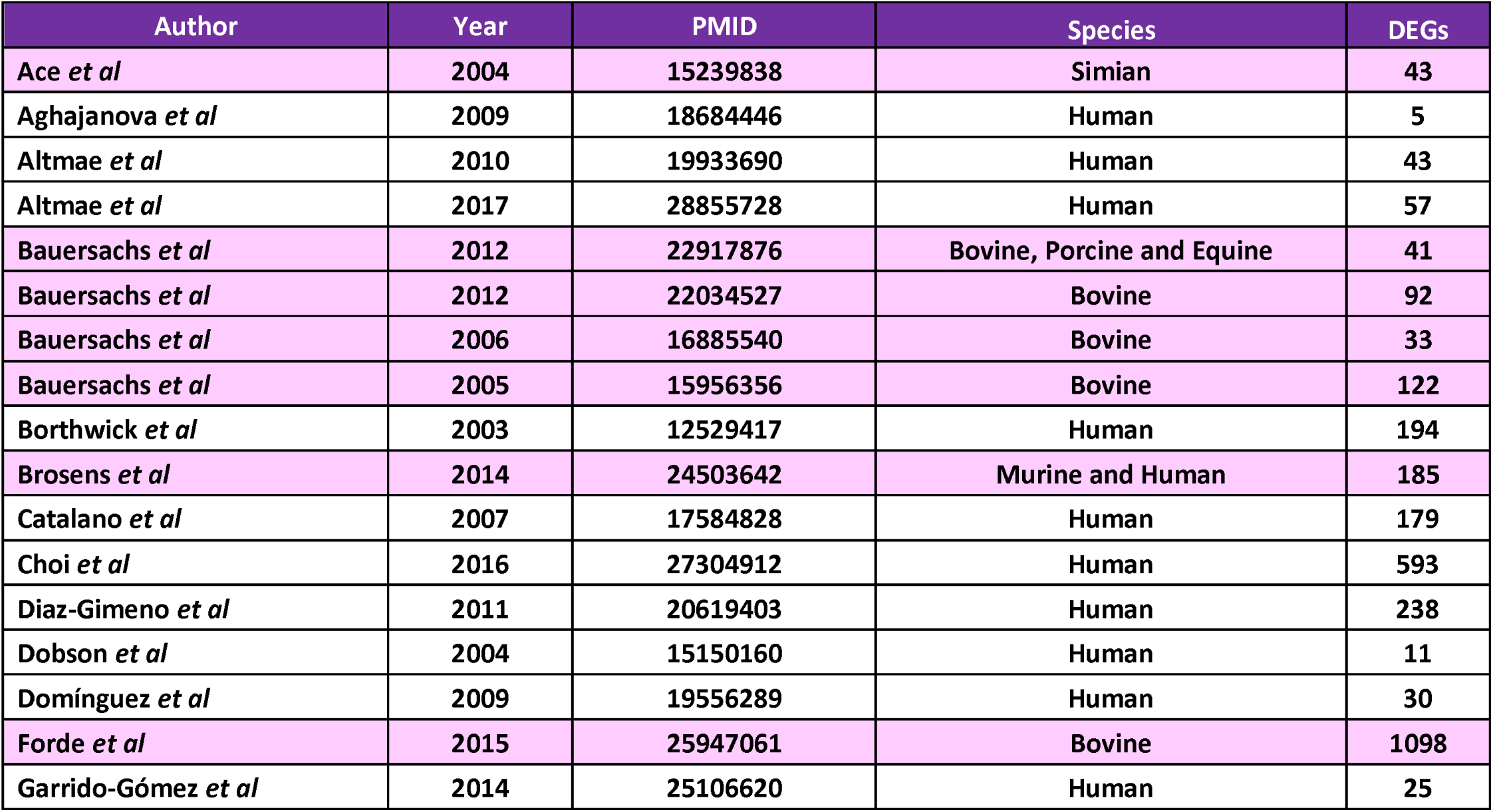
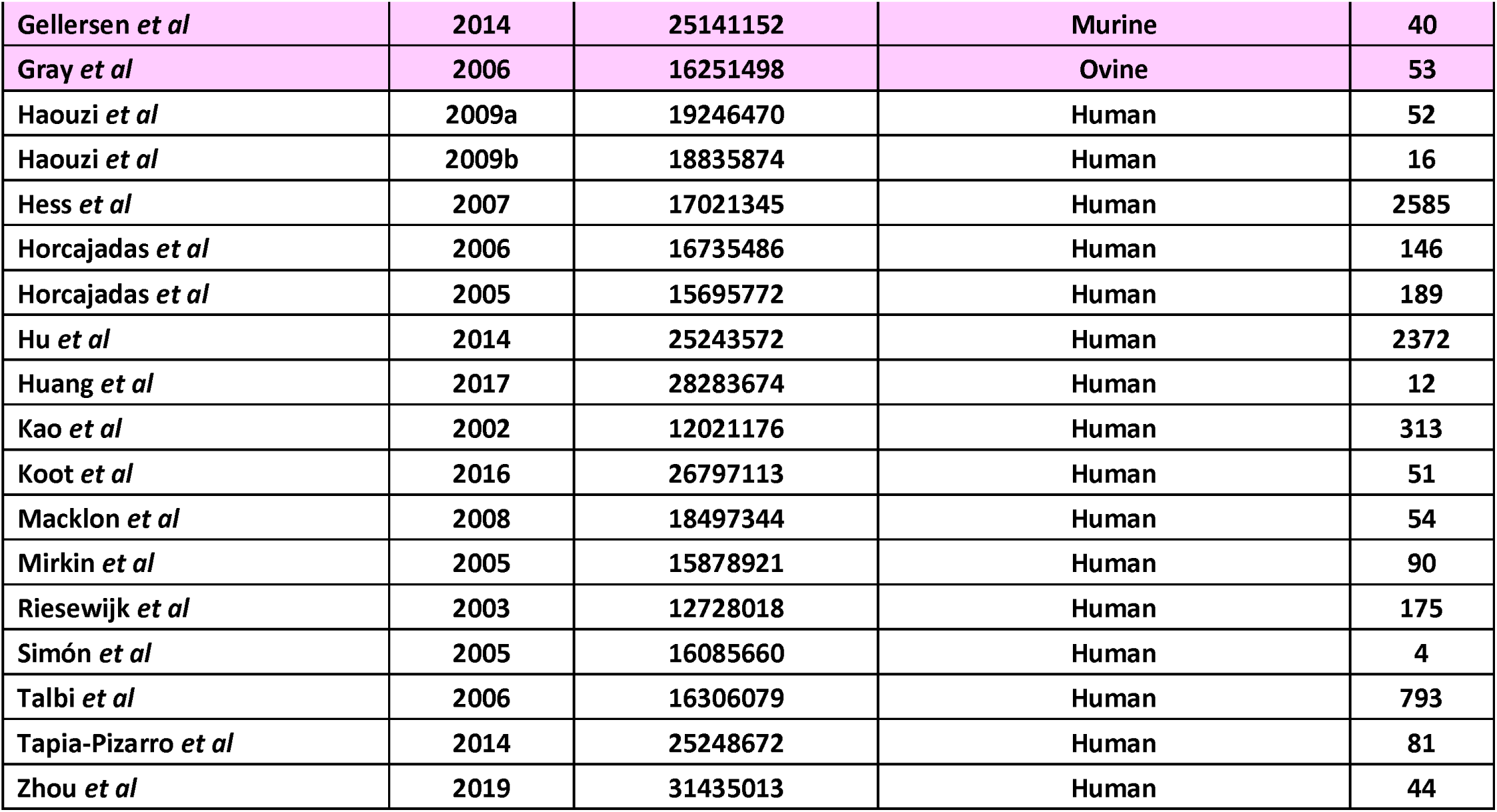
List of 35 studies published between 2002 and 2019 reporting endometrial DEGs (identified from PubMed literature searches)(9, 12, 14, 19, 36, 57, 102-130). From the pooled list of DEGs identified across all studies (duplicates removed), comparisons were made to try to validate proteomic data with computationally predicted miR-185-5p targets. Targets were then explore further during functional enrichment analyses with WebGestalt and g:Profiler. Non-human studies are highlighted in pink.

### Functional enrichment analysis with online toolsets

Two online toolkits were used to evaluate the pathways, functions and processes that are regulated by miR-185-5p. Functional enrichment analysis was initially performed using WebGestalt (WEB-based Gene SeT AnaLysis Toolkit; http://www.webgestalt.org/) (Wang et al., 2013). Protein lists for miR-185-5p mimic and inhibitor (or both) were analysed through WebGestalt and generated a list of the top 10 most significantly enriched gene ontology (GO) terms for biological processes (BP), molecular functions (MF), cellular components (CC), and KEGG pathways (Kanehisa, 2000). To ensure robustness in our analysis, g:Profiler (https://biit.cs.ut.ee/gprofiler/gost) generated a list of the mostly significantly represented GO terms, which were compared against those generated from WebGestalt.

### Pooling of targets and creation of ‘Proteins of Interest’ list

Following comparison of functional enrichment analyses between WebGestalt and g:Profiler, the most significantly overrepresented genes (within GO terms) were pooled and compared against published and predicted miR-185-5p targets, creating a miR-185-5p target list. Our criteria for “proteins of interest” were that proteins identified (from this study) whose corresponding gene was in the top 25% of the most frequently represented genes within GO terms. From this pooled list (Figure 1), those gene targets identified within shared function (e.g., protein binding genes) identified in miR-185-5p mimic and inhibitor targets) were selected as proteins of interest. STRING (https://string-db.org/) was used to determine protein-protein interaction networks of these miR-185-5p protein partners.

### Examination of target proteins in endometrial biopsies from women with RIF

#### Sample collection

Human luteal-phase endometrial biopsy samples were obtained from patients with recurrent implantation failure (n=10) and fertile controls (patients with a history of a previous clinical pregnancies or live birth; n=9). Ethical approval (18/WA/0356) was provided by the Tommy’s Biobank Tissue Access Committee and written informed consent provided by participants. Endometrial samples were obtained (using Wallach Endocell™ endometrial sampler) at the Implantation Research Clinic at University Hospitals Coventry and Warwickshire (UHCW), during the predicted pre-ovulatory LH surge in non-hormonally stimulated menstrual cycles. Overt endometrial pathology was excluded by transvaginal ultrasound prior to endometrial sampling. Patient characteristics are provided in Table 2.

**Table 2.**
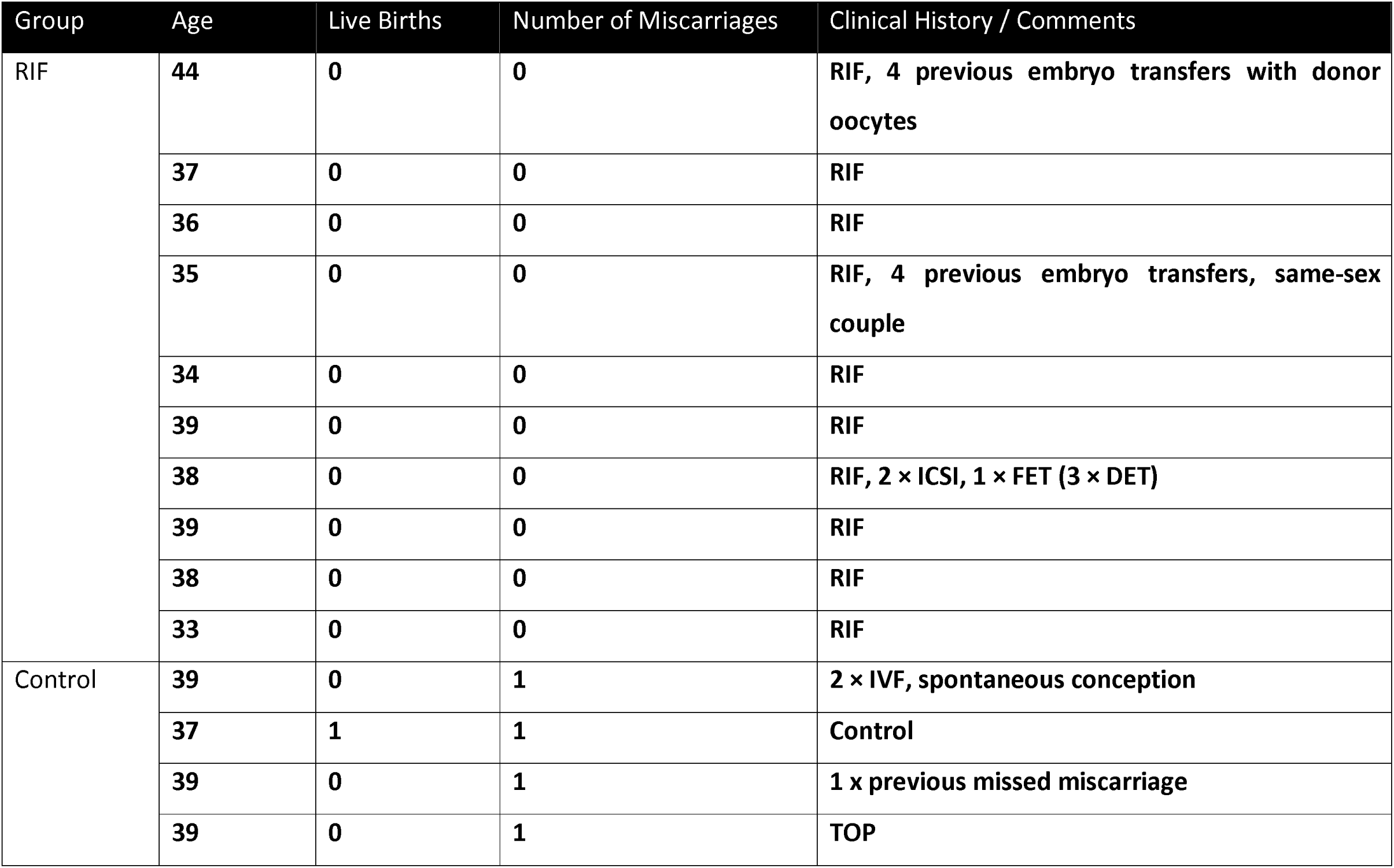
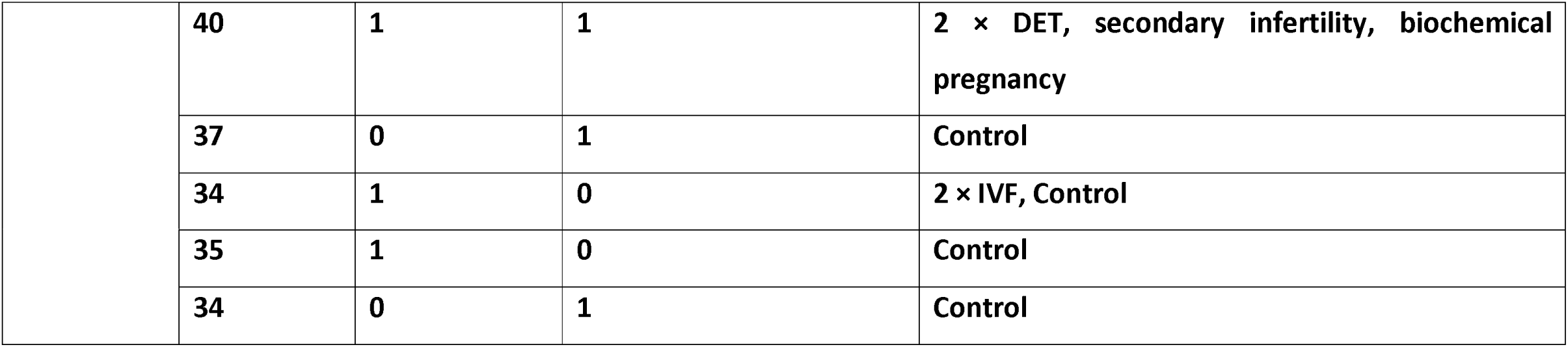
Summary of patient characteristics for which endometrial samples were recovered. RNA from endometrial biopsies from individuals with RIF (N=10) and fertile controls (N=10) who received unstimulated and timed mid-luteal endometrial biopsies, from which total RNA was extracted and used during analysis of cadherin binding protein target mRNAs.

#### RT-qPCR analysis of candidate genes

Total RNA was extracted using the RNeasy plus Universal Mini Kit (Qiagen®) as per manufacturer’s instructions and concentration determined using a NanoDrop-2000 spectrophotometer. Primers were designed using the online primer-basic local alignment search tool (BLAST) at NCBI (https://www.ncbi.nlm.nih.gov/tools/primer-blast/) (Table 3). Two hundred nanograms of RNA was reverse transcribed using a high-capacity reverse transcription kit (ThermoFisher Scientific) as per manufacturer’s instructions. Next, qPCR was carried out using Roche® LightCycler®96, using 5µL Roche® SYBR™ Green PCR Master Mix, 0.25μL of each (20μM) forward and reverse corresponding primer pair, and 2.5μL nuclease-free water and a final concentration 20ng of cDNA. The thermal cycling conditions were: 1) 95°C for 5 mins, 2) 45-cycles of 95°C for 10 secs, 56°C for 20 secs, and 72°C for 30 secs, 3) a melting curve analysis (95°C for 5 secs, rapid cooling to 65°C for 1 min, followed by a further slow increase in temperature to 97°C with fluorescence analysed with each 1°C increment). Relative expression values were calculated from raw Cq values and then normalised against the geometric mean of *ACTB* and *GAPDH* and differences determined using t-test with differences identified when p <0.05. A subset of these were evaluated further by exploring their protein-protein interaction networks using the STRING online database (https://string-db.org/) to identify interacting proteins.

**Table 3.**
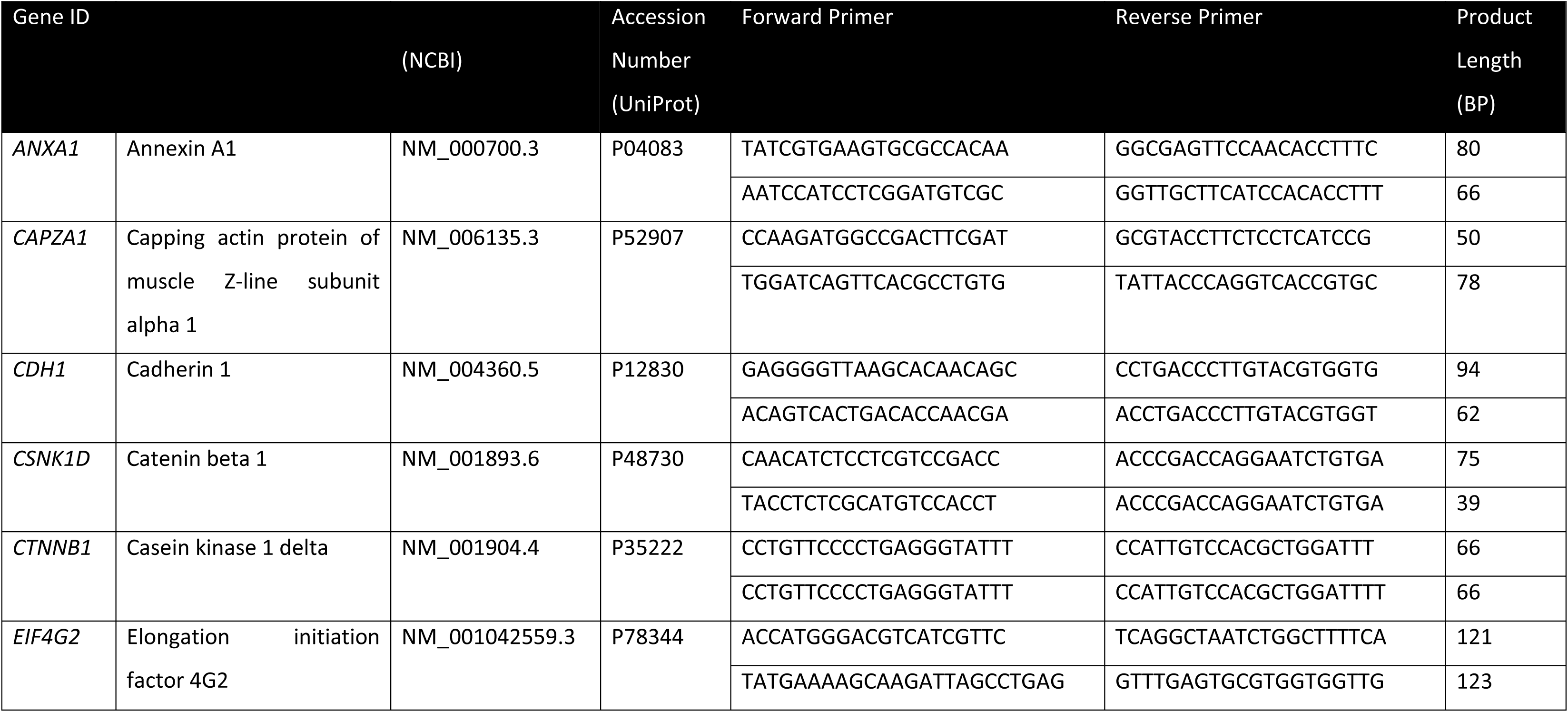
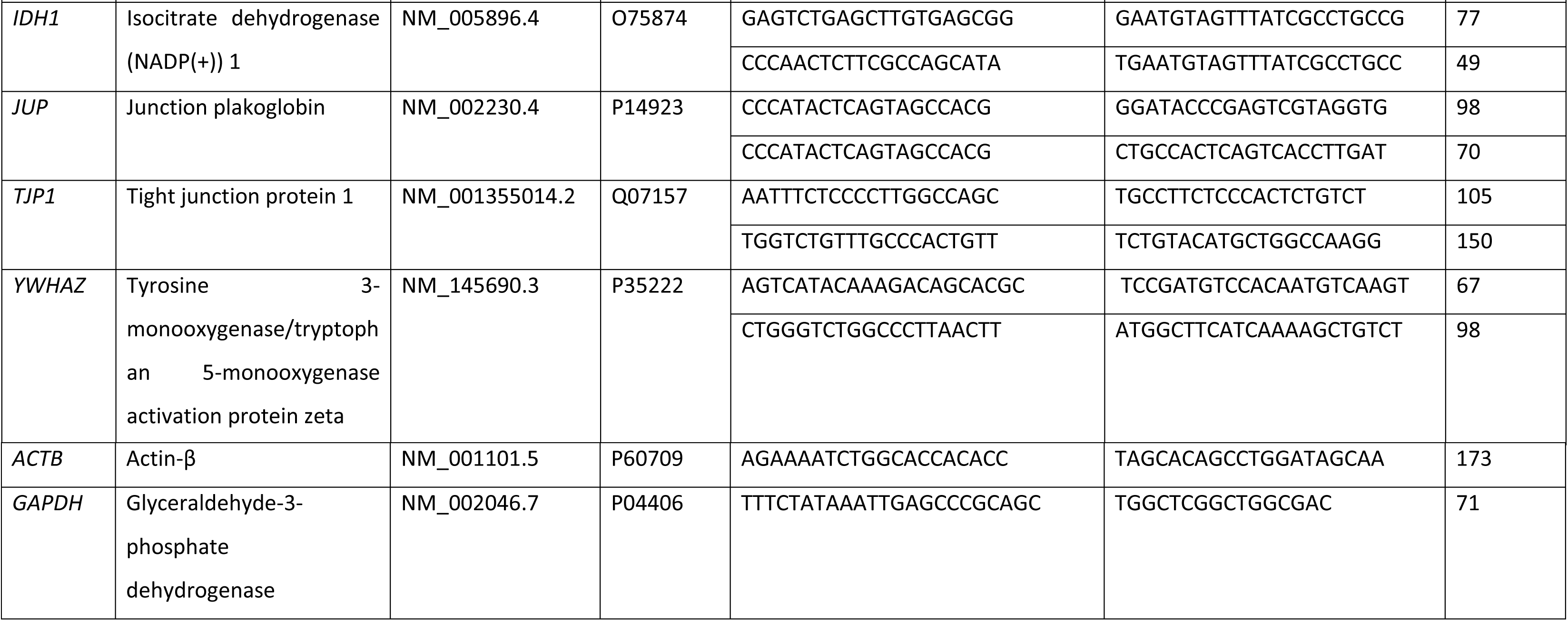
Primer designed for members of the cadherin binding protein (CBP) family. Gene. ID, descriptor, Refseq and UniProt accessions numbers for forward and reverse primers as well as product length. Primers were designed using NCBI BLAST and designed to span exon boundaries where possible.

## RESULTS

### miR-185-5p alters the endometrial proteome and implantation attachment

A significant difference in the ability of spheroids to attach to epithelia cells was observed between miR-185-5p mimic and inhibitor treated cells (p = 0.024), with a mean decrease in attachment (89.5%) following treatment with miR-185-5p mimic and increased attachment (96.8%) following treatment with miR-185-5p inhibitor (Figure 2A).

**Figure 2.**
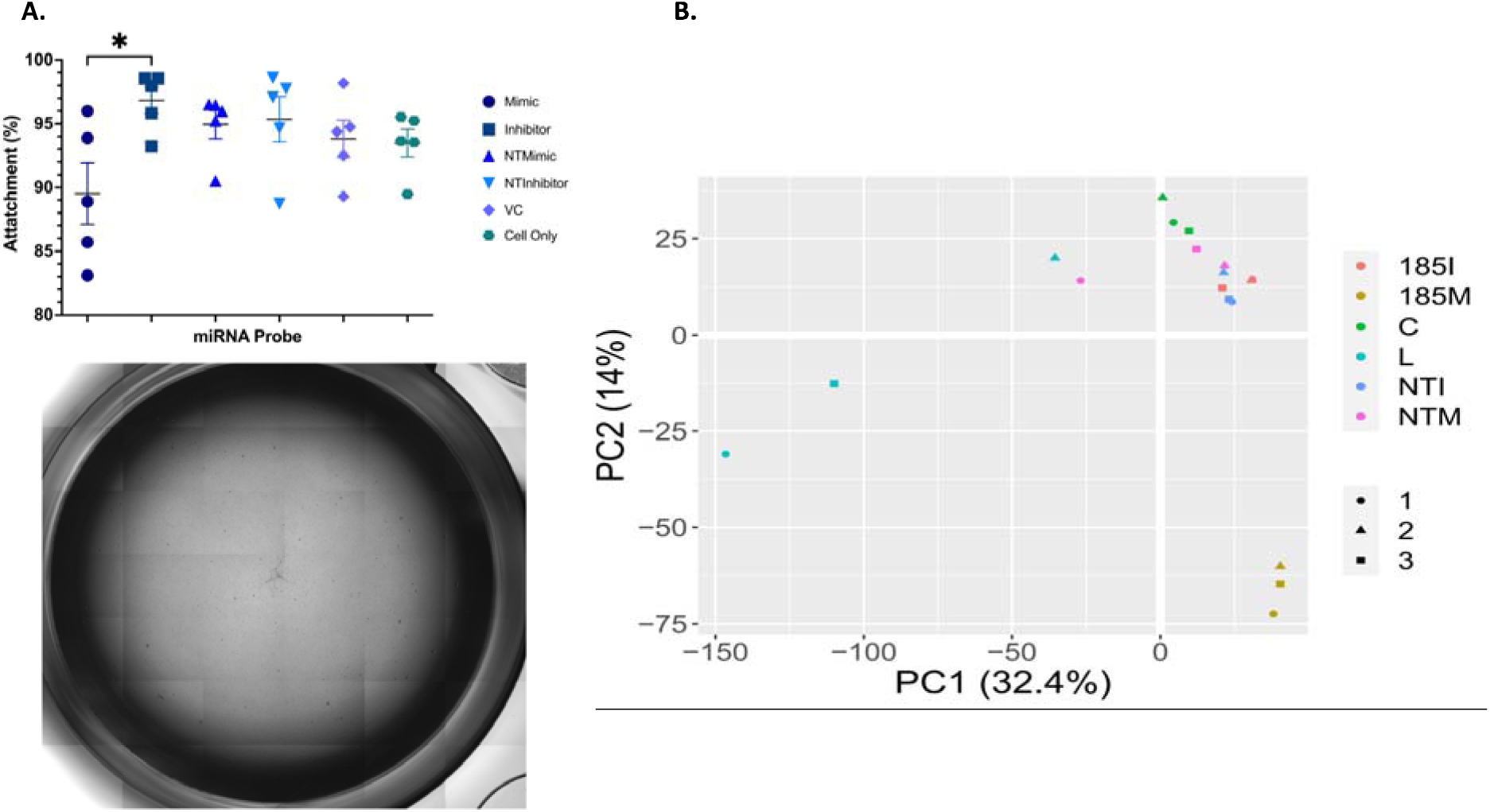
Impact of miR185 on endometrial epithelial cells. **A.** Implantation of spheroids in endometrial epithelia cells following treatment with miR-185-5p mimic, inhibitor, or recombinant protein PDI. Differences in spheroid-epithelial attachment percentage in response to miR-185-5p mimic or inhibitor transfection. Treatment groups include: C (Cell only); VC (Vehicle Control – DharmaFECT); NTM – non-targeting mimic; NTI – non-targeting inhibitor; Mimic miR-185-5p mimic; Inhibitor – miR-185-5p inhibitor. Differences in attachment percentage during pair-wise comparisons, specifically in miR-185-5p mimic versus miR-185-5p inhibitor-transfected hEECs are denoted by an asterix (p <0.05; unpaired two-tailed *t-*test). **B.** Principal Component Analysis (PCA) plot for proteomic analysis of endometrial epithelial cells following treatment with miR-185-5p mimic and inhibitor for 48 hrs. Samples were evaluated via TMT-mass spectrometry. in 3 biological replicates with sample IDs: C – Control (antibiotic-free media – DMEM/F12); L – Lipofectamine-2000 / also referred to as VC – vehicle control (lipofectamine-2000 + OptiMEM media); NTM – non-targeting mimic (miRIDIAN non-targeting mimic with lipofectamine-2000 and OptiMEM media); NTI – non-targeting inhibitor (miRIDIAN non-targeting inhibitor with lipofectamine-2000 and OptiMEM media); 185-M – miR-185-5p mimic (miRIDIAN miR-185-5p mimic with lipofectamine-2000 and OptiMEM media); 185-I – miR-185-5p inhibitor (miRIDIAN miR-185-5p inhibitor with lipofectamine-2000 and OptiMEM media).

Principal component analysis (PCA) plot demonstrated clear separation between controls and miR-185-5p mimic-treated samples, whereas inhibitor and non-targeting controls cluster with vehicle control (Figure 2B). To determine which proteins were changed in response to miR-185-5p mimic or inhibitor, a Venn diagram analysis was carried out to remove any off-target effects (Figure 3: All proteomic data are provided in Supplementary Table 1). Off target effects, *i.e.,* those changed in controls were removed from these lists (Figure 3), and a final list of proteins resulted in 1304 significantly changed in response to miR-185-5p mimic alone, 363 significantly changed is response to miR-185-5p inhibitor alone and 146 altered in response to both miR-185-5p mimic *and* inhibitor treatment (Figure 4).

**Figure 3.**
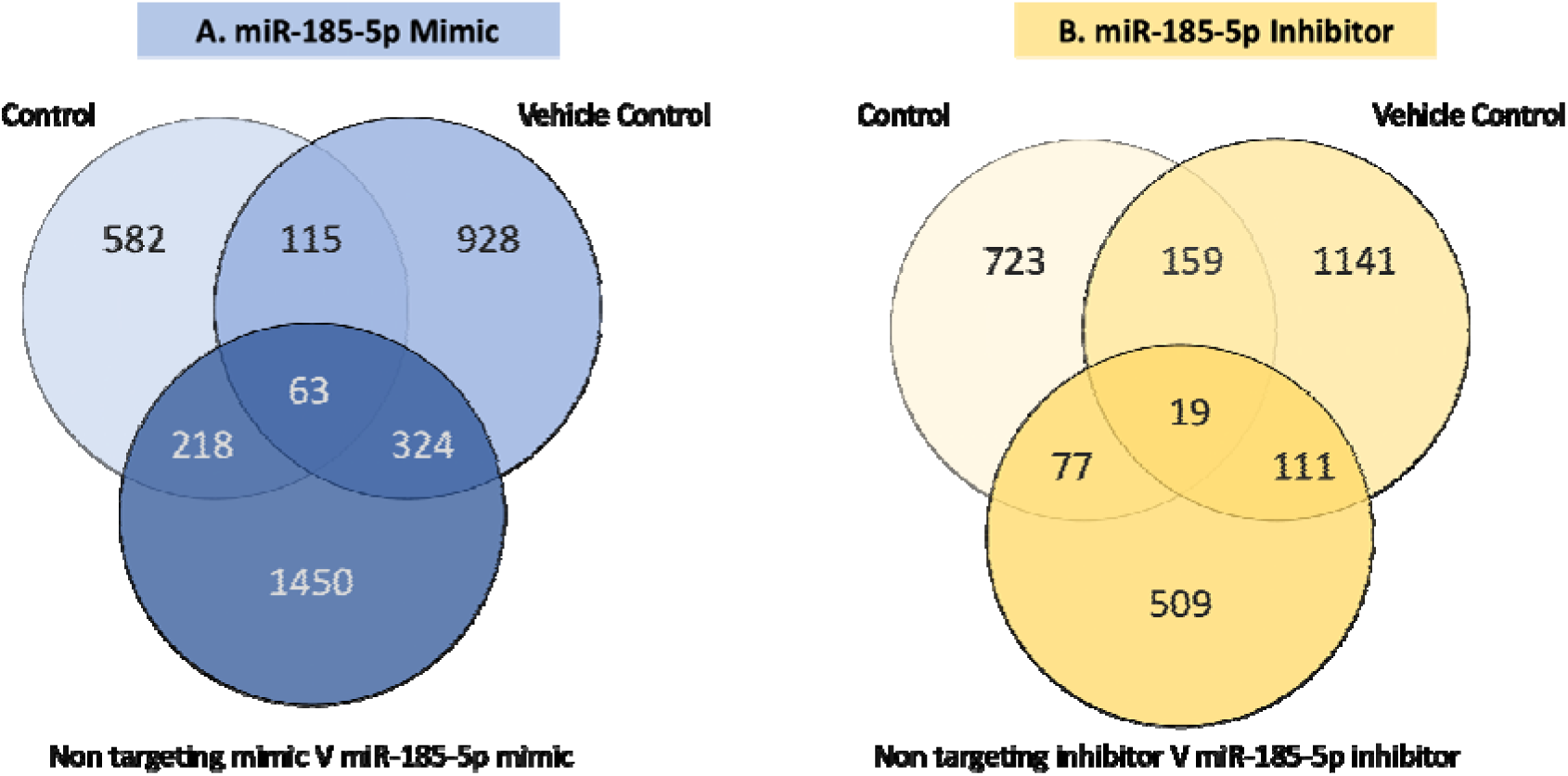
Venn diagram comparison of protein expression between treatment groups. To determine proteins that were altered specifically due to the actions of miR-185-5p different venn diagram analysis (created using Venny 2.1 https://bioinfogp.cnb.csic.es/tools/venny/) was used to remove those that were also changed in non-targeting mimic (NTM), non-targeting inhibitor (NTI), control (C) and vehicle control (VC) creating a pooled NTM/NTI versus control response and pooled NTM/NTI versus vehicle control response. These resulted in proteins only identified in response to miR-185-5p mimic (185-M: Left hand side) or inhibitor (185-I; Right hand side).

**Figure 4.**
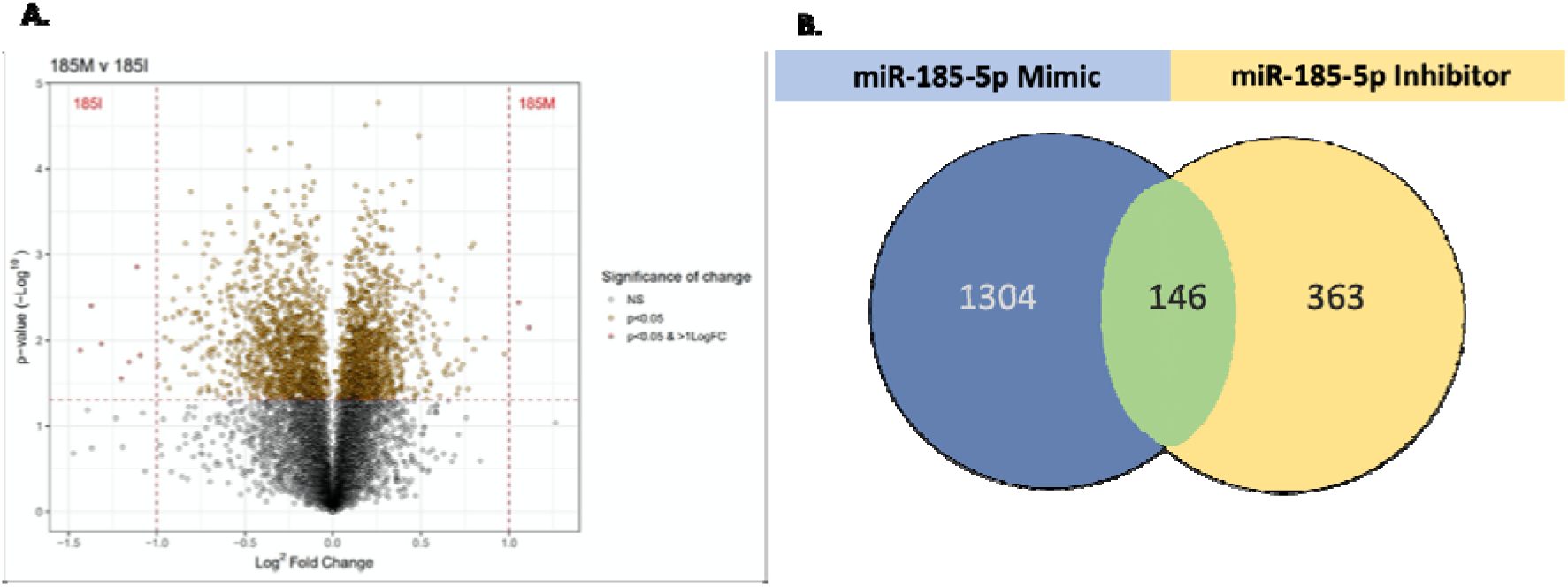
Proteins changed in endometrial epithelial cells in response to miR-185 treatment. **(A)** Volcano plot comparing miR-185-5p mimic-treated cells versus miR-185-5p inhibitor-treated cells, illustrating statistically significant log2FC (differentially expressed proteins) between miR-185-5p mimic and inhibitor samples. Proteins with the most statistically significant Log2FC (proteins that appear most abundant in response to treatment) are identified in the upper outer quadrants (red dots). Orange dots reveal proteins with a statistically significant difference in protein abundance (when comparing response between the two treatment groups), although with a log2FC less than 1 (less than a doubling in protein abundance) and therefore evaluating the functional relevance of these differences requires further scrutiny. **(B)** Identifying miR-185-5p mimic and inhibitor-specific targets and shared targets. Following identification of non-targeting mimic (NTM) and non-targeting inhibitor (NTI) protein response compared to control and vehicle control (Figure 3.9), protein targets specific to miR-185-5p mimic (185-M)-treated cells and miR-185-5p inhibitor (185-I)-treated cells could be identified. Venn diagrams were created with Venny 2.1 (https://bioinfogp.cnb.csic.es/tools/venny/). Cross comparison of these targets was performed to identify those specific to miR-185-5p mimic (n=1304) and miR-185-5p inhibitor (n=363). The intersection (n=146) represents those proteins identified in response to miR-185-5p mimic and inhibitor.

**Figure 5.**
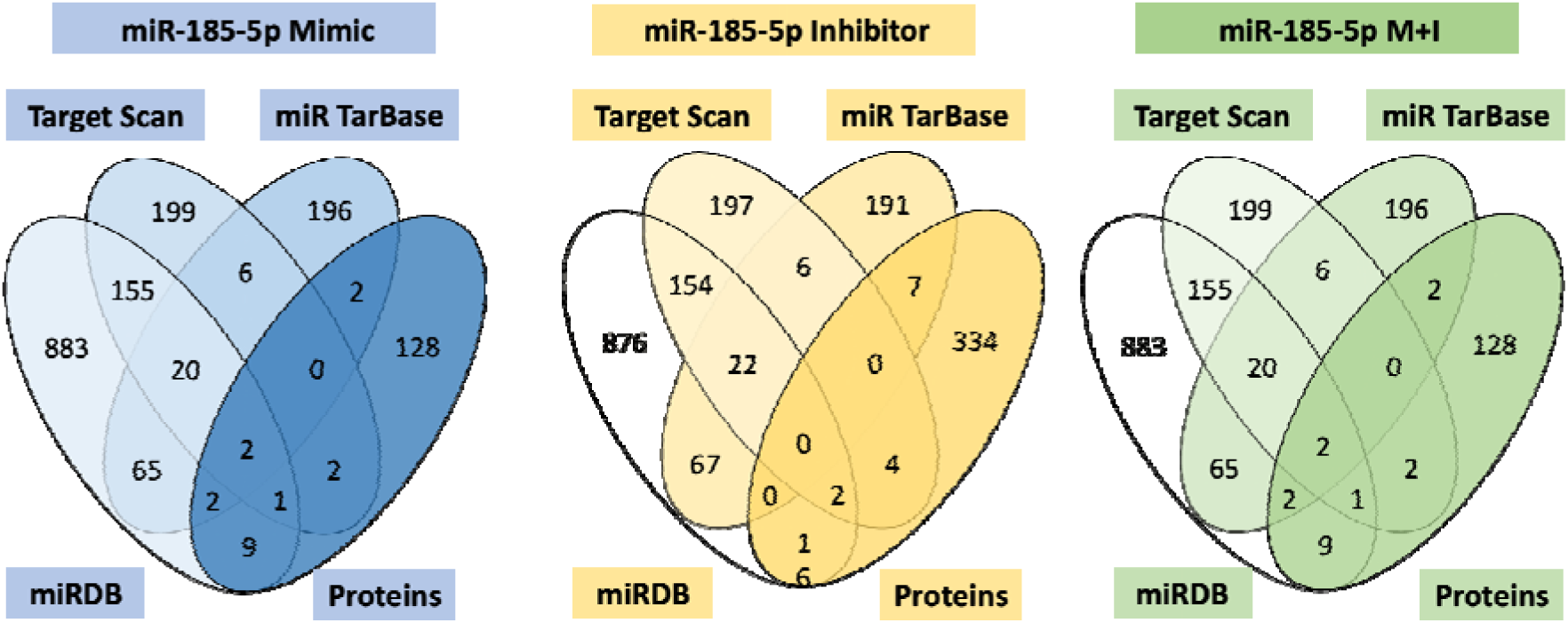
Venn diagrams illustrating validation of proteomics target data by direct comparison with each individual online miR-185-5p target prediction list. Venn diagrams shown are for mimic (miR-185-5p-M), inhibitor (miR-185-5p-I) and mimic *and* inhibitor (miR-185-5p-M+I) targets compared against miRDB, miRTarBase and TargetScan. Within the blue oval on each Venn diagram are the computationally predicted miR-185-5p targets also identified during miR-185-5p proteomic analysis. A large number of targets are shown not to be predicted by these three online databases. For example, 1151 miR-185-5p mimic targets were not predicted (although a number of these are published differentially expressed endometrial genes, as shown in the following section) and therefore may potentially be regulated indirectly. Venn diagrams were created with Venny 2.1 (https://bioinfogp.cnb.csic.es/tools/venny/).

### Functional Enrichment Analyses (Gene Ontology) for miR-185-5p targets

Over-represented biological processes, molecular functions, and cell components and KEGG pathways are provided in Supplementary Tables 2-5. Of particular interest were the targets for miR-185-5p that were found to be highly enriched for functions in cadherin binding and cell adhesion molecule binding. Similarly, independent analysis of overrepresented molecular functions for the targets of miR-185-5p using gProflier also identified enrichment of cadherin binding and protein binding (Supplementary Table 6).

As proteins identified in relation to cadherin binding, cell adhesion molecule binding and protein binding were identified as highly represented functions in miR-185-5p targets a pooled list of proteins from these three groups was created (Table 4) to highlight potential interactions between mimic (n=169), inhibitor (n=35), and mimic *and* inhibitor targets (n=22). The shared roles were identified between miR-185-5p mimic and inhibitor targets within the following BPs, MFs and CCs: cadherin binding (WG + gP), catalytic activity (gP), cell adhesion molecule binding (WG), cellular macromolecule metabolic process (gP), cytoplasm (gP), cytosol (gP), mitochondrion (WG), protein binding (gP), and RNA binding (WG).

**Table 4.** List of cadherin binding proteins used for target analysis. Members of cadherin binding proteins identified from miR-185-5p target prediction, published differentially expressed endometrial genes and functional enrichment analyses of proteomic analysis data. Ten were targets identified (shown in pink) for evaluation in RIF clinical samples with rationale given for each.

| miR-185-5p Cadherin Binding Targets |  |  |  |  |  |
| --- | --- | --- | --- | --- | --- |
| Mimic |  | Inhibitor |  | Mimic + Inhibitor |  |
| Predicted | Non-Predicted | Predicted | Non-Predicted | Predicted | Non-Predicted |
| CAPZA1 | CTNNB1 | CDH1 | ANXA1 | TJP1 | EIF2A |
| CSNK1D | EGFR | YWHAZ | EEF1G |  | EIF4G2 |
| EEF2 | EHD1 |  | RACK1 |  | HSPA8 |
| PPME1 | EPHA2 |  | TWF1 |  | IDH1 |
| RDX | JUP |  | YWHAE |  |  |
|  | KRT18 |  |  |  |  |
|  | RAB1A |  |  |  |  |
|  | RANBP1 |  |  |  |  |
|  | RANGAP1 |  |  |  |  |
|  | SH3GLB1 |  |  |  |  |

The pooled lists of common targets within cadherin binding, cell adhesion molecule binding and protein binding were cross compared with highly represented targets within GO terms (top 25%) and published and predicted miR-185-5p targets. These comparisons are shown in Figure 6 revealing key targets for miR-185-5p mimic (n=15), inhibitor (n=7), and both (n=5), including a number of both predicted and non-predicted targets, of which 10 of the most highly represented targets were selected for further evaluation in endometrial samples from RIF patients (Table 3).

**Figure 6.**
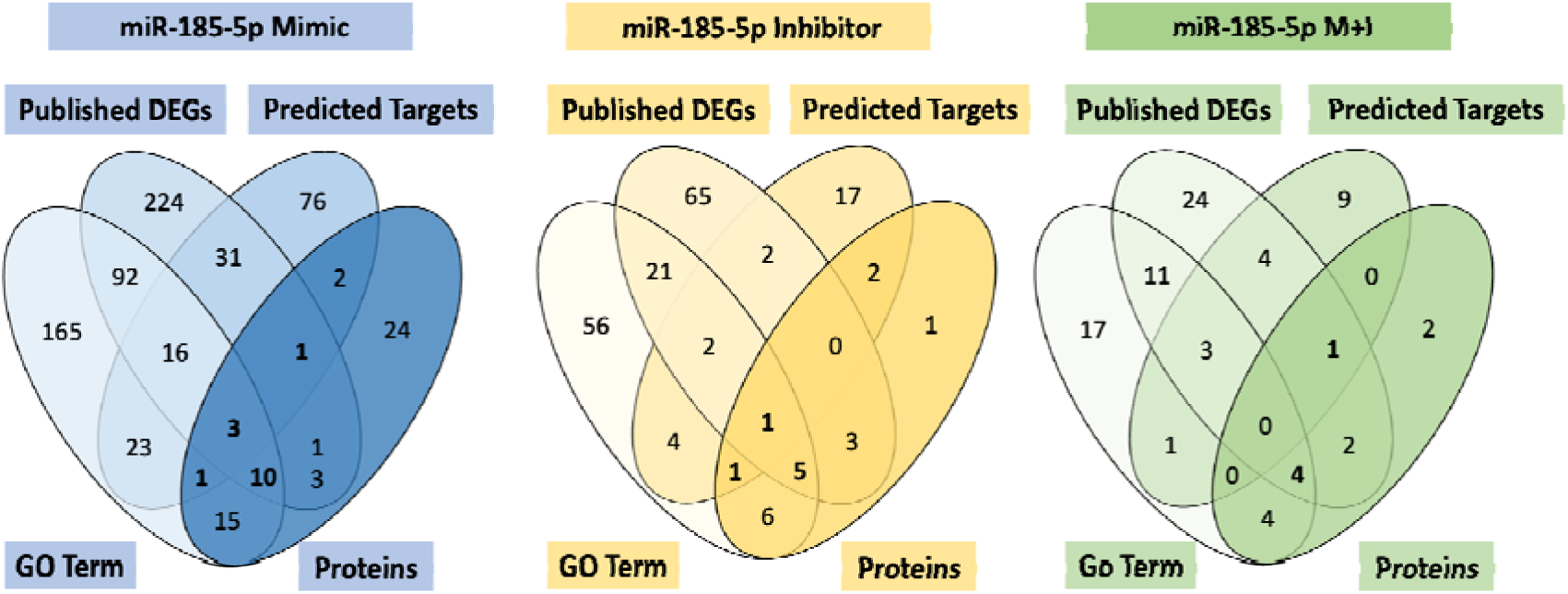
Cadherin Binding protein candidates. Venn diagrams illustrating significantly represented cadherin binding (CadBind) proteins (identified within pooled list from WG and gP; Table 3.21) and compared with published (PubDEG), predicted (PredT) and highly represented (Top25%GO) targets within GO terms (with Venn diagrams for each target group – miR-185-5p mimic [185-M], inhibitor [185-I] or both [185-M+I]). Venn diagrams were created with Venny 2.1 (https://bioinfogp.cnb.csic.es/tools/venny)

#### Published Endometrial DEGs and Relevance of miR-185-5p Targets

To determine whether protein targets of miR-185-5p that we identified in this study had been shown in other previously published data sets related to endometrial function, we compared our data with 35 studies published between 2002 and 2019 (Table 1). From these studies, 1557 DEGs were identified in the endometrium relating to functional and dysfunctional roles (Supplementary Table 7). Of these, 538 genes had protein targets that were identified in our proteomic data from this study, including targets for miR-185-5p mimic (n=390), miR-185-5p inhibitor (n=99) and both (n=49). Initially, only DEGs identified in 3 or more studies were to be used for subsequent cross-comparisons with proteomic data and predicted miR-185-5p targets. However, when evaluating proteins of interest later, this was deemed to be too stringent and excluded a large number of published DEGs. Therefore, any of the published targets were at least considered during early cross-comparisons.

### miR-185-5p Targets and Published Cyclic DEGs in the Endometrium

A previous study published by Talbi *et al* (2006) identified DEGs involved in proliferative (PE), early secretory (ESE), mid-secretory (MSE) and late secretory (LSE) endometrium (12). These data were interrogated further, and 793 DEGs identified as modified between different stages of the menstrual cycle. A comparison of these menstrual cycle-related changes in gene expression (from Talbi et al., 2006) with proteins altered by miR-185-5p in endometrial epithelia (our study; Table 4, and Supplementary Table 8) identified 12 targets in common, including cysteine and glycine rich protein 2 (CSRP2), decay-accelerating factor (DAF; also known as CD55), insulin-like growth factor binding protein 3 (IGFBP3), junction plakoglobin (JUP), nucleoporin 153 (NUP153), solute carrier family 7 member 1 (SLC7A1), solute carrier family 25 member 1 (SLC25A1), serine peptidase inhibitor, Kunitz type 1 (SPINT1) and squalene epoxidase (SQLE). Three miR-185-5p inhibitor targets were also identified, including cadherin 1 (CDH1), lysophospholipase 1 (LYPLA1) and transglutaminase 2 (TGM2). Specifically, CSRP2, NUP153, SLC25A1, SLC7A1 and SQLE were shown to be downregulated, whilst IGFBP3 and TGM2 were upregulated in the proliferative to early secretory phase endometrium. In contrast, CDH1, DAF, JUP and SPINT1 were shown to be downregulated in the mid-secretory to late-secretory endometrium.

### STRING Interactions for miR-185-5p Targets

From the overall list of miR-185-5p pooled targets (cadherin binding, cell adhesion molecule binding and protein binding) including mimic (n=169), inhibitor (n=35) and both mimic *and* inhibitor (n=22), protein-protein interactions (PPIs) were evaluated (on 03/06/2022) for each group to determine the strength of association between each miR-185-5p target using STRING DB (Szklarczyk et al., 2019). Protein-protein interaction (PPI) enrichment values were statistically significant for miR-185-5p mimic targets (p value < 1.0e^-16^) and miR-185-5p inhibitor targets (p value = 5.91e^-7^) but not statistically significant for miR-185-5p mimic *and* inhibitor. Nevertheless, cadherin binding and cell adhesion molecule binding remained significantly represented among these mimic *and* inhibitor targets (FDR of 0.0300 and 0.0048, respectively). Cadherin binding, cell adhesion molecule binding, protein binding among miR-185-5p mimic targets had an FDR of 1.29e^-05^, 0.00089, and 4.75e^-17^, respectively. For miR-185-5p inhibitor targets, cadherin binding and protein binding were shown to have an FDR of 0.0022 and 0.0002, respectively. Figure 8 demonstrates PPIs in the pooled highly represented miR-185-5p targets (cadherin binding, cell adhesion molecule binding and protein binding), with cadherin binding targets specifically highlighted in red.

#### Expression of CBP mRNA for CBP members in endometria from women experiencing RIF

All members of the CBP mRNAs investigated were detected in biopsies from women irrespective of their fertility status (Figure 7), except for *EIF4G2* which could not be amplified reliably. Statistical analysis of CBP mRNA expression revealed an increase in expression of two members of the cadherin binding family of transcripts, with significantly reduced (absolute) expression values for Casein kinase 1 isoform delta (*CSNK1D:* p = 0.0363) and a borderline significant reduction in (absolute) expression for *TJP1* (p = 0.0788).

**Figure 7.**
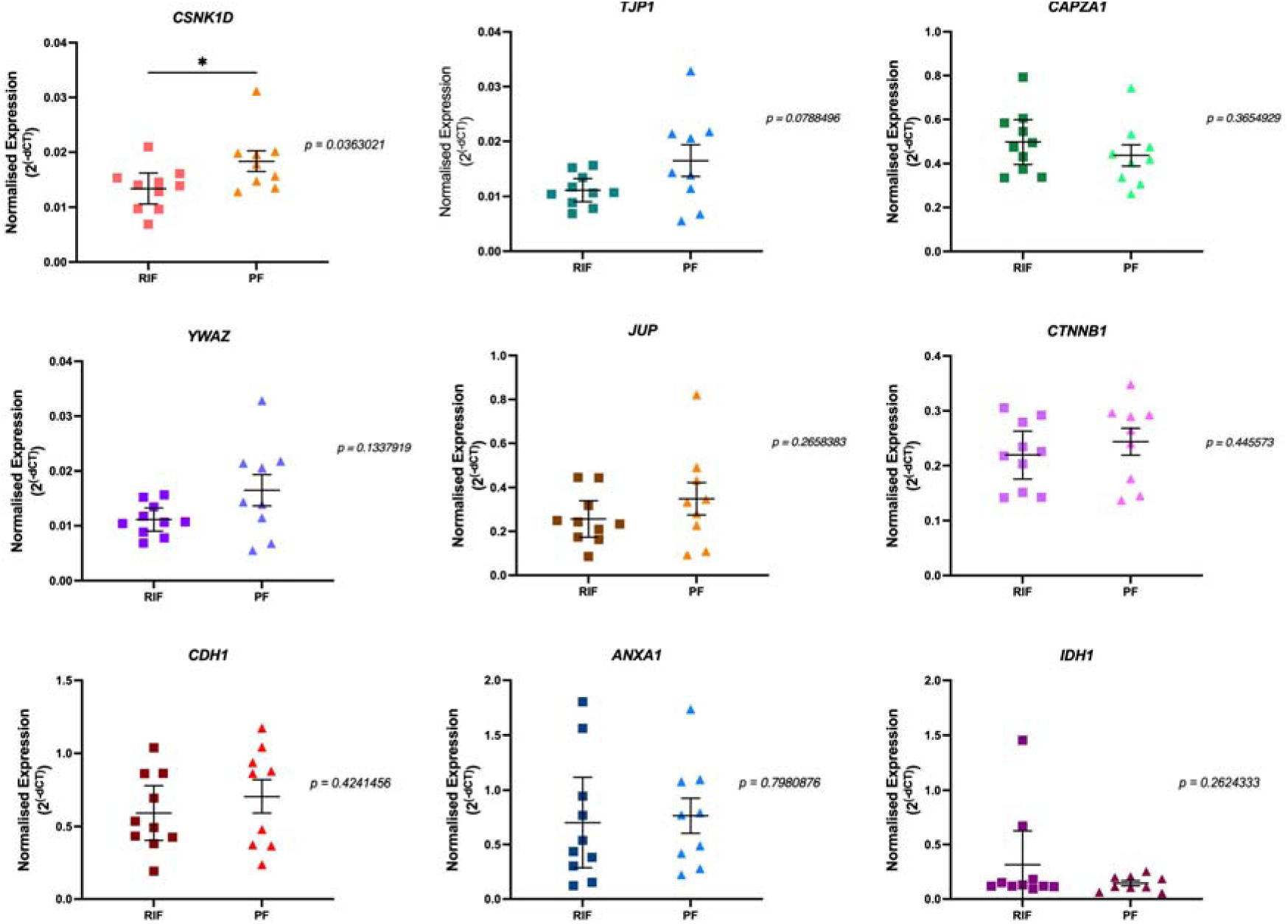
Expression of members of the CNP family in endometria from women with and without RIF. Normalised expression (2^-ΔCq^) for selected CBP mRNAs in RIF (squares) and fertile controls (triangles; PF) normalised expression values (2^-ΔCq.^) +/-SEM (n=10 and n=9, for RIF and successful pregnancy groups, respectively) and displayed as a measure of the spread of expression values across each clinical group. An Asterix. (*) depicts a. significant difference in expression value when p < 0.05.

Protein-protein interaction network analysis (Figure 8) showed CSNK1D has significant interactions with cryptochrome circadian regulators CRY1/CRY2 *and* period circadian regulators PER1/PER2, p53 (*TP53*) and Bystin (*BYSL*). Significantly enriched biological processes for CSNK1D including the circadian regulation of gene expression (p = 3.21e^-9^), negative regulation of intracellular steroid hormone receptor signalling pathway (p = 3.62e^-7^), regulation of cell communication (p = 6.18e^-5^) and regulation of cell differentiation (p = 0.00024). Significantly enriched molecular functions include beta-catenin binding (p = 1.51e^-6^), transcription factor binding (p = 0.00045) and E-box binding (p = 0.0056). The PPI network for CSNK1D is also shown to be highly represented in KEGG pathways for circadian rhythm (p = 1.80e^-12^), Wnt signalling (p = 0.00010), adherens junctions (p = 0.0486) and several cancer pathways, including transcriptional misregulation in cancer (p = 0.0239).

**Figure 8.**
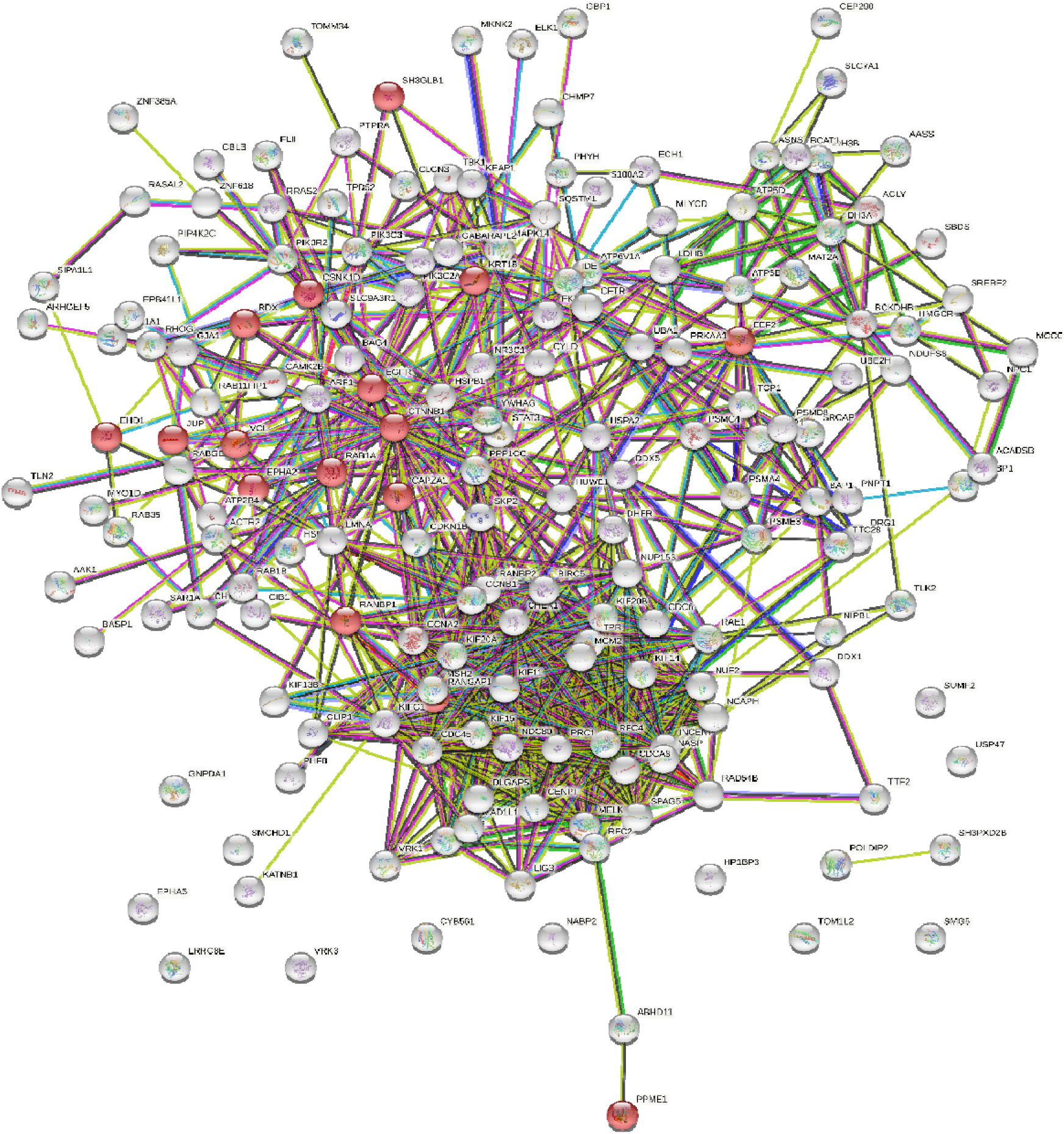

TJP1 was shown to have significant interactions with claudin-2 (CLDN2), catenin-β-1 (CTNNB1), junctional adhesion molecule-1 or F11-receptor (F11R), junctional adhesion molecule-3 (JAM3), gap junction protein alpha-1 (GJA1), and proto-oncogene tyrosine-protein kinase Src (SRC). The PPI enrichment for TJP1 interactions is also statistically significant (p = 2.6e^-12^). Significantly enriched biological processes included cell-cell junction organisation (p = 1.34e^-17^), adherens junction organisation (p = 8.87e^-12^), maintenance of blood-brain barrier (p = 3.7e^-9^), apical junction assembly (p = 1.22e^-7^), establishment of endothelial barrier (p = 2.78e^-5^), regulation of canonical Wnt signalling (p = 0.0018) and regulation of vascular permeability (p = 0.0027). Molecular functions significantly enriched include beta-catenin binding (p = 5.39e^-15^), cadherin binding (p = 5.51e^-8^) and cell adhesion molecule binding (p = 7.82e^-8^). The PPI network for TJP1 is also highly represented in KEGG pathways for adherens junctions (p = 3.08e^-10^), tight junctions (p = 6.34e^-10^), Wnt signalling (p = 2.63e^-8^), leucocyte transendothelial migration (p = 1.52e^-7^), cell adhesion molecules (p = 4.54e^-7^), signalling pathways regulating pluripotency of stem cells (p = 0.0067), focal adhesion (p = 0.0126) and hypoxia-inducible factor-1 (HIF-1) signalling pathways (p = 0.0482).

Unsurprisingly, given the association (and proximity) of the epithelial cadherins and catenins, catenin-β-1 significantly interacts with *both* CSNK1D and TJP1. However, there are a number of other second-shell interactors that have been shown to be related to both CSNK1D and TJP1 (identified using STRING DB (Szklarczyk et al., 2019)), including tumour suppressor adenomatous polyposis coli (APC), vinculin (VCL) and s-phase kinase associated protein 1 (SKP1). These proteins are also involved in relevant biological processes, including beta-catenin destruction complex assembly (APC; p = 2.08e^-11^), adherens junction assembly (VCL; p = 0.0092), and SCF-(SKP1-CUL1-F-box protein) ubiquitin ligase complex assembly (SKP1; p = 6.46e^-6^).

## DISCUSSION

We sought to understand if an evolutionarily conserved microRNA – miR-185-5p – and the targets it regulates in the endometrium may contribute to recurrent implantation failure in humans. Our data demonstrate that modification of endometrial epithelial cells *in vitro* with mimics or inhibitors of miR-185-5p alters the ability of the epithelia to facilitate attachment of BeWo spheroids. Proteomic analysis of endometrial epithelial cells following transfection with mimic and inhibitors of miR-185-5p demonstrated that evolutionarily conserved miR-185-5p alters a large number of proteins in pathways relevant to implantation events. Although not all computationally predicted as miR-185-5p targets, a large list of published endometrial DEGs (n=538) suggests that miR-185-5p has wide-reaching interactions with pathways that could potentially become dysregulated during implantation. Two specific components of the cadherin binding family of transcripts were found to be differentially expressed in luteal phase endometrial biopsies from RIF patients, demonstrating an important role for miR-185-5p and its downstream targets in successful implantation.

This proteomic study has allowed the comparison of miR-185-5p targets with previously published data sets. The list of genes generated will serve as a reference point for evaluating other targets, which may become clinically apparent in future studies, irrespective of their computational target prediction at this time. Those targets computationally predicted but not identified in current proteomic datasets may reflect cell culture conditions and may be worth re-considering in future experiments. Given that many were not computationally predicted targets of miR-185-5p, it may be that miR-185-5p also affects a wide range of mechanisms related to implantation indirectly. This is supported by the identification of common roles for miR-185-5p targets (shared across miR-185-5p targets in response to mimic, inhibitor or both), including cadherin binding, cell adhesion molecular binding and protein binding, all of which are relevant for early embryo apposition. It is also noteworthy that metabolic pathways were also highly represented among miR-185-5p mimic targets in KEGG pathways, emphasising their potential regulatory role during the increased metabolic demands associated with early implantation events (RoyChoudhury et al., 2016). Of the shared roles identified for miR-185-5p targets, the most significant function identified was cadherin binding. Cadherin binding has been shown to be highly represented amongst miR-185-5p predicted, published and functionally-enriched targets (including those related to other relevant pathways, including cell adhesion molecule binding and protein binding). The effects of miR-185-5p on implantation attachment mechanisms has been explored with an implantation model (implantation assay) and validated cadherin binding targets have been evaluated in RIF clinical samples to identify any significant differences between patients with RIF and fertile controls (proven fertility; PF).

From the 10 candidate transcripts associated with CBP, we evaluated expression of nine of these genes. *CSNK1D* and *TJP1* were identified as differentially expressed, in biopsies from women who experienced RIF. Casein kinase 1 isoform delta (CSNK1D) is part of a group of evolutionarily conserved serine/threonine kinases with a multitude of functions, including those related to cell-cycle progression (and tumour progression), apoptosis, circadian rhythm, vesicle trafficking and Wnt-signalling (Peters et al., 1999; Behrend et al., 2000; McKay et al., 2001; Stöter et al., 2005; Bernatik et al., 2011; Mazzoldi et al., 2019), functions with relevance to endometrial function and early pregnancy events. Inhibition of casein kinase 1 isoforms have been shown to alter cytoplasmic domain E-cadherin-phosphorylation, decreasing cell-cell adhesive activity *in vitro* (Dupre-Crochet et al., 2007). CSNK1D specifically has also been shown to regulate PER2 through GAPVD1 (GTPase Activating Protein and VPS9 Domains 1), affecting downstream functions within the mammalian circadian clock (Ibrahim et al., 2021). Interestingly, PER2 has also been associated with synchronization of endometrial proliferation and aperiodic decidual gene expression (Muter et al., 2015). It also has a regulatory role in tumorigenesis by enhancing erythropoietin production in tumour cells under hypoxic conditions by phosphorylating hypoxia-inducible factor-2 (HIF-2α) (Pangou et al., 2016); which may be a potential survival strategy utilised during restrictive early embryonic histotrophic nutrition. Taken together, these various roles for CSNK1D highlight its potential functional significance during implantation, with emphasis on peripheral (endometrial) clock functions which are likely to be relevant during cyclic decidualisation.

Tight junction protein 1 (TJP1), also known as zonula occludens 1 (ZO-1), is a member of a group of transmembrane scaffold proteins whose primary role is to regulate luminal epithelial contacts with adjacent cells, forming continuous belt-like structures (through multiple different interactions) (Heinemann and Schuetz, 2019). Loss of *TJP1* has been shown to alter human trophoblast cell-cell fusion and differentiation (Pidoux et al., 2010; Dunk et al., 2012). *TJP1* has also been shown to be co-ordinately expressed with epithelial cadherin (E-cadherin) within stromal cells of the primary decidual zone (PDZ) and appears to have a role in the development of semi-permeable barrier functions within the decidua which enable nutritive support for the invading embryo, whilst also shielding it from potentially hostile maternal antigenic immune responses (Paria et al., 1999). Dissolution of tight-junctions, decreased claudin and occludin expression, and remodelling of the extracellular matrix are fundamental features of epithelial-mesenchymal transition (EMT), a process which facilitates embryo invasion through the maternal luminal (endometrial) epithelium (Huang et al., 2012; Lamouille et al., 2014). EMT is also associated with altered desmosome activity and decreased levels of connexin (Yilmaz and Christofori, 2009; Bax et al., 2011; Huang et al., 2012), contributing further to disruption of cell-cell adhesion mechanisms. In addition, tight junctions are also essential during the maintenance of endothelial permeability barriers (Tornavaca et al., 2015), with ZO-1 deficient mice having been shown to have a lethal phenotype with abnormal embryonic vasculature identified during yolk sac angiogenesis (Katsuno et al., 2008). A recent study of vasculature remodelling in bladder tumours identified TJP1 as being highly and consistently expressed in invasive tumours and correlated with tumour angiogenesis, in keeping with TJP1 upregulation of C-C motif chemokine ligand 2 (CCL2) via twist family bHLH transcription factor 1 (TWIST1) (Liu et al., 2022). These roles suggest loss of TJP1 function may significantly impair implantation in patients with RIF.

In addition to CSNK1D and TJP1, other significant proteins of their interacting networks, have already been implicated in human embryo implantation and reproductive failure, although the majority of these interacting proteins were not identified in our proteomic data or panel of published endometrial DEGs, highlighting the potential for indirect effects of miR-185-5p on other implantation-related proteins.

Bystin (*BYSL*) is a soluble cytoplasmic protein which forms a complex with tastin (*TROAP*) and trophinin (*TRO*) to regulate cell adhesion between trophoblast and the endometrial luminal epithelium, with TRO being highly dependent on BYSL and TROAP in order to have functional cell adhesive activity (Fukuda et al., 1995; Suzuki et al., 1998). Interestingly, although embryos from trophinin-null mice do not survive, all three proteins disappear from placental villi by 10 weeks gestational age (Suzuki et al., 1999), highlighting their significant yet temporal role during early embryo attachment and invasion.

Claudin-2 (CLDN2) is key member of the claudin family of tight junction proteins. CLDN2 specifically, is a cation-selective channel forming tight junction protein which has been identified as a feature of leaky epithelia (Furuse et al., 2001) and has been more recently identified in biological processes such as proliferation, migration and cell-fate decisions (Venugopal et al., 2019). Recently CLDN2-containing EVs were identified as a biomarker for managing colorectal cancer liver metastasis (Tabariès et al., 2021).

Tight junctions and adherens junctions constitute different cell-to-cell contacts with discrete functions, such as the regulation of ions and small molecules and maintenance of apical-basal polarity, or the regulation of cell-to-cell adhesion, respectively. Cingulin (CGN) and occludin (OCLN) associate with claudins and ZO-proteins in the formation of tight junctions. Vinculin (VCN), an actin binding protein, associates with the β-catenin-plakoglobin-P120-cadherin complex in adherens junctions (Dejana et al., 2009).

Gap junction protein alpha 1 (also known as connexin-43) is a recognised transmembrane protein which regulates intercellular communication and is essential during decidualisation. Inhibition of connexin-43 in mice leads to abnormal decidualisation and loss of appropriate vascular expansion to maintain pregnancy (Ramathal et al., 2010). Intriguingly, expression of connexin-43 appears to be specifically upregulated in endometrial stroma at day 11-15 of the menstrual cycle, with subsequent loss of expression during the luteal phase (Jahn et al., 1995). Interestingly, IL-1β, a recognised embryonic signalling cytokine (Krüssel et al., 2003) has been shown to inhibit connexin-43, by halting decidualisation via the extracellular signal-regulated kinase (ERK) 1/2 signal transduction pathway and has been implicated in decidual dysfunction in patients with endometriosis and infertility (Yu et al., 2019).

S-phase kinase associated protein 1 (SKP1) is a member of the SCF (SKP1, Cullin-1, F-box containing) complex of E3 ubiquitin ligases that target substrates with poly-ubiquitin chains, in preparation for proteasomal degradation. The SCF-complex is made up of 69 SCF-E3 ubiquitin ligases, although the specific function of each remains unclear (Thompson et al., 2021). Via a p53-induced pathway, SKP1-interactions have been shown to reduce the activity of (or destruction of) beta-catenin and directly inhibit beta-catenin-dependent T-cell factor (TCF) and lymphoid enhancer factor (LEF) transcription factors (Matsuzawa and Reed, 2001); the major end point of Wnt signalling and responsible for cell fate decisions (Cadigan and Waterman, 2012). More recently, Cullin-3 (CUL3), another scaffold protein, also responsible for the assembly of ubiquitin ligase complexes, like SCF complexes, has been shown to be highly expressed in VCT and EVT and may also be involved decidualisation, with CUL3 small interfering RNA (siRNA) found to reduce MMP activity via TIMP1 and TIMP2, and CUL3 knockdown mice found to have less invasive trophoblast cells (Zhang et al., 2015). It is possible therefore, that dysregulation of the SCF pathway might also have similar functions relating to preparation of the endometrial stroma for implantation.

The impact of evolutionarily conserved miR-185-5p on wider implantation mechanisms appears irrefutable. Our application of a very simple model of early implantation has provided clear evidence of miR-185-5p regulating mechanisms which alter trophoblast-epithelial attachment. Within the limitations of an epithelial-cell-only attachment model, miR-185-5p appeared to be anti-adhesive in its effect on spheroid-epithelial attachment. However, given that miR-185-5p targets are differentially expressed in RIF patient biopsies, it is entirely possible that miR-185-5p has anti-adhesive effects at the level of the luminal epithelium, whilst coordinating pathways within the stromal compartment which facilitate invasion beyond the epithelium; mechanisms that could not be altered in this implantation assay model, warranting further investigation. *CSNK1D*, *TJP1* and their interacting network proteins appear to have highly relevant functions to decidualisation and implantation, and further evaluation is recommended. Finally, investigating altered function(s) of these CBP targets and their interacting proteins *in vitro* (and eventually *in vivo*) will help to establish the clinical impact of these regulators and proteins in RIF; effects that may also be of relevance in patients with recurrent early pregnancy loss.

## ACKNOWLEDGEMENTS

Research in NF’s lab was supported by N8 agri-food pump priming, QR GCRF, as well as BBSRC grant numbers BB/R017522/1, and BB/X007332/1 and Wellcome Trust grant 227178/Z/23/Z. We thank the people who donated tissue.

